# SoS Notebook: An Interactive Multi-Language Data Analysis Environment

**DOI:** 10.1101/274001

**Authors:** Bo Peng, Gao Wang, Jun Ma, Man Chong Leong, Chris Wakefield, Melott James, Yulun Chiu, Di Du, John N. Weinstein

## Abstract

**Motivation:** Complex bioinformatic data analysis workflows involving multiple scripts in different languages can be difficult to consolidate, share, and reproduce. An environment that streamlines the entire processes of data collection, analysis, visualization and reporting of such multi-language analyses is currently lacking.

**Results:** We developed Script of Scripts (SoS) Notebook, a web-based notebook environment that allows the use of multiple scripting language in a single notebook, with data flowing freely within and across languages. SoS Notebook enables researchers to perform sophisticated bioinformatic analysis using the most suitable tools for different parts of the workflow, without the limitations of a particular language or complications of cross-language communications.

**Availability:** SoS Notebook is hosted at http://vatlab.github.io/SoS/ and is distributed under a BSD license.

## 1 Introduction

Due to complications and limitations of cross-language interfaces (e.g. the R and MATLAB interfaces for Python), most bioinformaticians write separate scripts when tools in different languages are needed for parts of the analysis. Adding to all of the difficulties in interfacing between languages and producing reports from multiple scripts, it is challenging to keep track of separate script files, share fragmented workflows with colleagues, and archive the workflows properly for future reference.

A multi-language data analysis environment provides a unified platform on which workflows composed in mixed languages can be easily executed, shared, and reproduced. A few such environments under active development include RStudio Notebook, Apache Zeppelin, and Beaker Notebook (now BeakerX). RStudio has a clear root in R with limited support for other languages; Zeppelin is designed for data analytics and visualization in large-scale data exploration; Beaker Notebook has now been replaced by BeakerX which supports only Java-based languages. None of those environments support SAS or MATLAB, and that limits their applications in the bioinformatics and biostatistics community.

With the goal of developing a versatile environment for daily bioinformatic data analysis, we developed a multi-language notebook environment powered by a Python3-based workflow engine entitled Script of Scripts (SoS), hence its name SoS Notebook.

## 2 Features

SoS Notebook consists of a new kernel and a number of front-end extensions of the web-based Jupyter Notebook platform. The SoS kernel acts as a proxy to the SoS workflow engine and a hub between Jupyter and more than 60 existing Jupyter kernels. It executes scripts in multiple languages and coordinates data exchange among the kernels. The SoS front-end extends the single-kernel notebook interface of Jupyter to a multi-language notebook interface with global and cell-level language selectors and a multi-purpose side panel. Using a set of “magics” (special commands prefixed with %) and keyboard shortcuts, SoS Notebook provides an interactive notebook environment with explicit and automatic kernel switch and data exchange, line-by-line code execution, and other useful features such as preview of variables and files.

**Fig 1.**
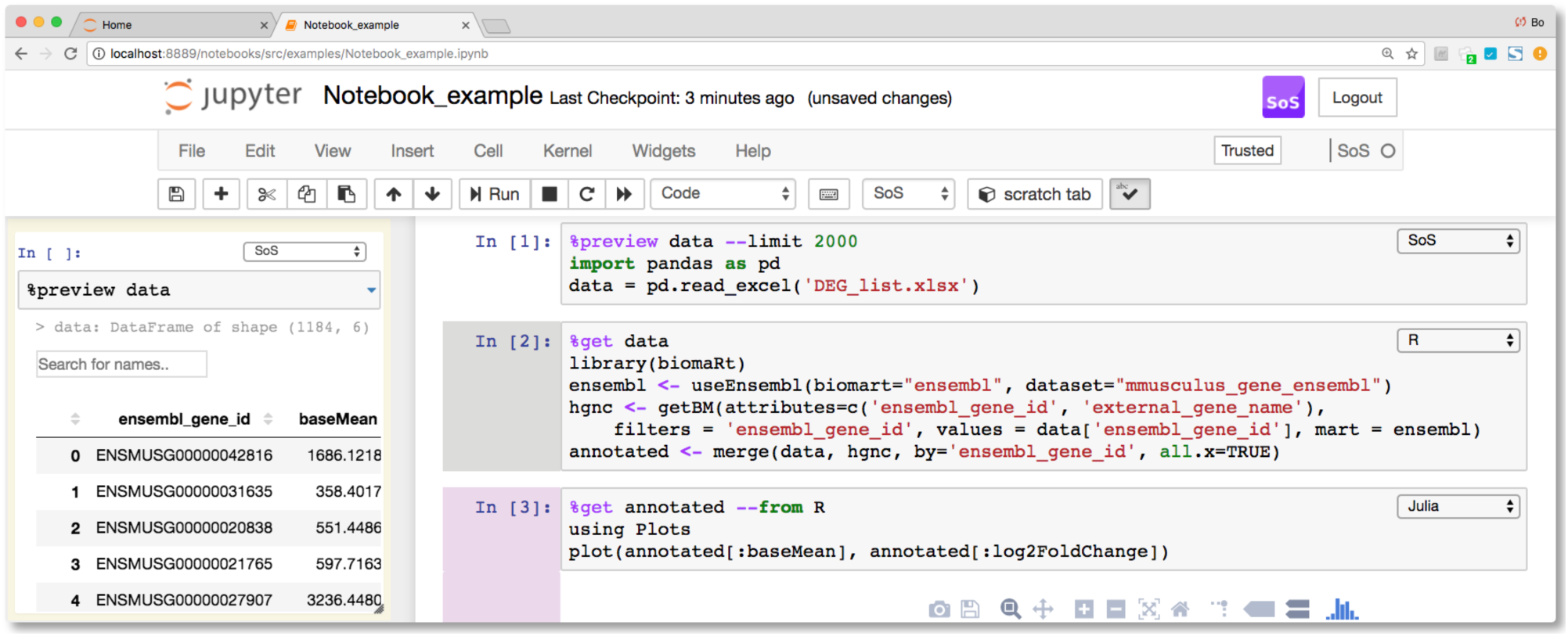
An example SoS notebook that uses SoS (Python) to read data from an excel file, preview it in the side panel as a searchable and sortable table, transfer data to R to annotate ensembl gene id with HGNC names using the Bioconductor package biomaRt, then plot the annotated data in Julia.

An SoS notebook consists of markdown cells and code cells with content executed by either SoS (based on Python 3) or one of the subkernels. Cell kernels can be changed either interactively or using magics `%with` and `%use`. Magic `%expand` interpolates cell contents with variables in the SoS kernel, allowing the composition of scripts in different languages using shared notebook variables (e.g. filenames, parameters). Magic `%capture` captures cell outputs as SoS variables so results from one kernel can be passed to and used by other kernels. Along with the main notebook, a side panel is provided to perform a variety of nonpermanent actions, such as executing scratch commands, showing results of line-by-line execution, previewing variables and files, and showing the table of contents of the notebook.

A more powerful data exchange method is provided to exchange variables among kernels for supported languages (Bash, JavaScript, Julia, MATLAB, Octave, Python2 and 3, R, and SAS). Because of large differences in datatypes among scripting languages, SoS transfers variables through the creation of independent homonymous variables of the most similar datatypes in the destination language. For example, although `3` and `c(3, 5)` are both numeric arrays in R, they are transferred to Python as integer 3 and numpy array([3, 5]) respectively. Similarly, Julia’s Char and str types are both converted to and from Python as str, while Python `DataFrame` is converted to `data.frame` in R, dataset in SAS, `table` in MATLAB, `dataframe` in Octave, and nested dictionaries in JavaScript. Data exchanges are performed automatically for variables with names starting with sos and explicitly with magics such as `%put` and `%get`. Interestingly, magic “`%with language --in v1 --out v2`” evaluates the contents of a cell in another language with input and output variables as if calling a function in the current kernel.

SoS magics provide functions for smooth, interactive data analysis. Of particular interest is a `%preview` magic that previews variables and expressions from any kernel, and local or remote files in many different formats, including bioinformatics-specific formats such as `fastq, vcf`, and `bam`. The preview magic could be triggered explicitly or automatically (e.g. during line-by-line execution) and can preview dataframes in scrollable, sortable, and searchable tabular format, or in interactive scatterplots with a tooltip for each data point. In addition, magic `%render` renders headers, tables, and figures from texts produced by any kernel, allowing the generation of a single report from results obtained using multiple languages.

SoS Notebook facilitates the generation of reports from multilanguage data analysis by providing magics to capture and render results from multiple kernels, and tags, shortcuts, and templates to generate reports in HTML format. For example, the `sos_report template` allows generation by default of HTML reports that display only selected material. A reader can therefore focus immediately on the core messages of a report and display hidden source code, text, and results if needed. Output from the `%preview` magic can be included to report large datasets as dynamic tables and plots.

## 3 Discussions

SoS Notebook provides an interactive working environment in which users can apply the most suitable languages and tools for different parts of a data analysis. Resulting notebooks contain detailed descriptions, complete source codes, and results. They can be readily shared with others and re-executed to reproduce prior analyses. SoS Notebook also serves as an execution and management console for the SoS workflow system, as described in a companion report (in preparation). This unique combination of interactive environment and workflow engine makes it easy to convert scripts developed for interactive data analysis to a workflow for large-scale data crunching, further distinguishing SoS Notebook from other multi-language environments. Although SoS currently focuses on commonly-used languages for bioinformatic applications under a Jupyter front end, we are working with the open source community, especially the JupyterLab (successor of Jupyter) team, to support more languages and to port SoS Notebook to JupyterLab.

## Acknowledgements

We appreciate feedback and support from members of the Department of Bioinformatics and Computational Biology at UT MD Anderson Cancer Center. We also appreciate reviewers and users of SoS Notebook for their very helpful feedback.

